# Language-associated brain representations covary with disruption of selfhood under psilocybin

**DOI:** 10.64898/2026.10.06.757069

**Authors:** Greg Cooper, Devon Stoliker, Vincent Sebastian Astl Delgado, George Blackburne, Leonardo Novelli, David W. Green, Ravi Das, Adeel Razi, Jeremy I. Skipper

**Affiliations:** Clinical Psychopharmacology Unit, Department of Clinical, Educational & Health Psychology, University College London, London, United Kingdom; Turner Institute for Brain and Mental Health, School of Psychological Sciences, Monash University, Melbourne, Victoria, Australia; Experimental Psychology, Division of Psychology and Language Sciences, University College London, London, United Kingdom; Department of Computing, Imperial College London, London, United Kingdom; Monash Biomedical Imaging, Monash University, Melbourne, Victoria, Australia; CIFAR Global Scholars Program, Toronto, Ontario, Canada

**Keywords:** psilocybin, selfhood, ego dissolution, language, naturalistic fMRI, representational similarity analysis, hyperalignment

## Abstract

Language helps organise experience into an idiosyncratic, temporally extended sense of self. We therefore used fMRI to investigate whether greater similarity between subjects in the activity of language-associated brain regions accompanied psilocybin-induced loss of selfhood using fMRI. Participants heard music and viewed moving clouds at baseline and after 19 mg psilocybin. At each brain location and time point, we tested whether similarity increased with each pair’s summed rank on the Ego Dissolution Inventory (EDI; N=43). We grouped independent components of these associations by time course and decoded each group’s map with Neurosynth. Elementary visual imagery (SI; N=49) and baseline activity served as comparators. Under psilocybin, one group formed a distributed, left-lateralised frontotemporal network matching the Neurosynth language map (r=.382, spatial-null p<.001, q=.003), nearly twice the largest comparator estimate (SI: r=.208, p=.004, q=.028; baseline: r≤.12). In exploratory analyses, resting-state connectivity of this network decreased with posterior visual cortex and increased with thalamic and frontal regions. Participants with higher EDI scores showed larger changes (ρ=.455, Bonferroni-adjusted p=.048; SI: ρ=.008), most strongly for items describing loss of ego or identity (ρ=.43–.51; other items, ρ=.22–.33). We propose that language-associated systems incorporate each person’s semantic and autobiographical context into experience, and that disrupting them contributes to loss of ego and identity.

## Introduction

> *“…I did not know that I am. I lived in a world that was a no-world. I cannot hope to describe adequately that unconscious, yet conscious time of nothingness. I did not know that I knew aught, or that I lived or acted or desired*.*”*
>
> — Helen Keller
>
> *“I had a nothing mind, a flotsam mind. I was incredibly focused on the present, with very little awareness or interest in my past or future. My entire environment felt interconnected, like cells in a large, breathing organism. To experience this Quiet was to be it*.*”*
>
> — Lauren Marks

A reader could be forgiven for taking either epigraph as an account of a high-dose psychedelic experience. The first passage is Helen Keller on the years before Anne Sullivan taught her language (Keller, 1908); the second is Lauren Marks, after a ruptured aneurysm left her with a left perisylvian lesion and severe expressive aphasia (Marks, 2017). Both describe ineffability, unity with the surroundings, loss of agency and dissolution of identity, closely paralleling constructs measured by widely used instruments of psychedelic phenomenology (Studerus et al., 2010; Barrett et al., 2015; Nour et al., 2016; Sleight et al., 2023). Comparable descriptions recur across accounts of language loss and recovery (Moss, 1972; Taylor, 2009), while studies of aphasic stroke document profound disruption of narrative identity (Shadden, 2005; Musser et al., 2015). The recurrence of this phenomenology across developmental language deprivation, acquired aphasia and psychedelic states may therefore reflect a common aetiology in the disruption of language-mediated representation of one’s self to one’s self.

Language contributes to this self-organisation from perception through reflective selfhood. Linguistic labels shape categorisation, perception and memory and can bring otherwise suppressed visual stimuli into awareness (Lupyan and Ward, 2013; Lupyan et al., 2020). Language also lets people take themselves as an object of thought and relate the present to a remembered past and anticipated future (Gallagher, 2000; Blanke and Metzinger, 2009; Christoff et al., 2011; Letheby, 2020; Millière, 2020). Inner speech internalises dialogue with others and allows a person to evaluate their own thoughts from the position of a generalised other (Mead, 1934; Morin, 2006; Alderson-Day and Fernyhough, 2015), while narrative and developmental accounts tie identity and autobiographical memory to socially scaffolded narration (Fivush and Nelson, 2004; McAdams and McLean, 2013).

On the other hand, the temporary loss of one’s sense of self (ego dissolution) while under the effects of psychedelic drugs has been studied primarily at the level of regional and large-scale network changes, rather than in terms of modulation of specific cognitive functions. Its intensity has been found to covary with reduced parahippocampal coupling and integrity of a salience-associated network under psilocybin (Lebedev et al., 2015), increased global connectivity of angular and insular cortex under LSD (Tagliazucchi et al., 2016), altered medial prefrontal glutamate (Mason et al., 2020), changes in directed coupling among default-mode-, salience- and attention-related systems (Stoliker et al., 2023, 2024) and stronger connectivity between dorsolateral prefrontal cortex and thalamus under LSD (Coleman et al., 2025), within a broader picture of cortical desynchronisation and a compressed unimodal-to-transmodal hierarchy (Girn et al., 2022; Siegel et al., 2024). These effects depend on context, weakening when participants perform a task or watch a video (Siegel et al., 2024; Mediano et al., 2024). In the PsiConnect cohort analysed here, psilocybin integrated networks that ordinarily segregate internal and external processing and aligned brain dynamics with experimental context, such that participants reporting deeper self- and boundary-dissolving experiences showed stronger alignment to context (Stoliker et al., 2026).

Language is one candidate for the cognitive process these changes disrupt: lacking it in development or losing it after stroke produces experiences much like ego dissolution, and its role in maintaining a felt sense of selfhood (categorising percepts, relating the present to past and future, and sustaining a narrative identity) is putatively suspended by ego dissolution (Millière, 2020; Stoliker et al., 2022). Indeed, altered language is among the oldest quantified effects of psychedelics, from reversible word-finding and expressive disturbances to reduced linguistic predictability and syntactic complexity (Stoll, 1947; Jarvik et al., 1955; Isbell, 1959; Amarel and Cheek, 1965; Landon and Fischer, 1970). Modern studies similarly show altered semantic processing: psilocybin increases indirect semantic priming, while LSD preserves naming speed but increases semantically related substitutions and alters the organisation and semantic structure of spontaneous speech (Spitzer et al., 1996; Family et al., 2016; Sanz et al., 2021; Wießner et al., 2023). Psilocybin also reduces the electrophysiological distinction between self-generated and externally altered vocal feedback, in proportion to changes in connectedness and meaning (Smigielski et al., 2020). Consistent with this behavioural literature, the cortical distribution of 5-HT2A receptors, the principal target mediating psychedelic effects, is spatially associated with meta-analytic Neurosynth maps for terms including semantic, language, words and reading (Preller et al., 2018; Tagliazucchi, 2022). From these findings we hypothesise that ego dissolution involves a transient disruption of language-mediated self-organisation, wherein the systems that ordinarily impose conceptual, temporal and personal structure on experience lose some of their usual influence.

How that should appear in the brain depends on the neurobiology of language assumed. On one view a small set of core perisylvian regions, principally the left inferior frontal gyrus (Broca’s area) and posterior superior temporal cortex, uses whatever context is available to predict upcoming input, and does so by dynamically recruiting a periphery of sensory, motor, mnemonic and affective regions (Skipper, 2015, 2022). Because the context a listener brings is their own accumulated history of associations, the anatomical configuration of the periphery recruited by a shared stimulus is dynamic and person-specific, and averaging across people retains only the cores they hold in common (Aliko et al., 2023; Drijvers et al., 2025). These cores are compatible with evidence for a reproducible and functionally selective left frontotemporal network (Fedorenko et al., 2024).

For selfhood, on this account, these cores provide sites at which a person’s stored semantic and autobiographical associations can be co-active with current percepts, linguistic or not (Skipper, 2022). Within the shared cores (Huth et al., 2016), fine-grained activity patterns still differ between people (Haxby et al., 2020), because each person brings different associations to the same input. Hence, if language-mediated self-organisation is disrupted, personal associations are less co-active with current percepts, so people reporting greater ego dissolution should show patterns within language cores that are less idiosyncratic.

This prediction can be tested directly with naturalistic stimulation, which gives every participant the same evolving sensory sequence while leaving their memories, expectations and interpretations free to differ, such that variability between people can be measured rather than averaged away (Finn et al., 2020). Because less idiosyncratic patterns are more similar between people, the similarity between two participants’ core patterns should increase with how strongly both report ego dissolution, measured here with the Ego Dissolution Inventory (EDI; Nour et al., 2016). Finn et al. (2020) call this an Anna Karenina model, after the opening line of Tolstoy’s novel: “Happy families are all alike; every unhappy family is unhappy in its own way” (Tolstoy, 1878/1901). The present application inverts the valence, with dissolved egos expected to resemble one another and intact ones to remain intact each in their own way.

We tested this in the PsiConnect cohort, in which participants completed identical music and silent cloud-viewing tasks at baseline and after 19 mg oral psilocybin (Novelli et al., 2026), using hyperalignment and time-resolved spatial inter-subject representational similarity analysis (SIS-RSA; Ma and Skipper, 2025) to relate the neural similarity of each participant pair to the Anna Karenina model (Figure 1). We predicted this association under psilocybin, in language-associated regions centred on inferior-frontal and temporal cortex and separate from the regions responding to the sounds and images themselves. We used elementary imagery as a phenomenological comparator, and related post-dose scores to baseline activity to test whether any language association reflected pre-existing variation. Because language shapes perception (Lupyan et al., 2020), we also asked, in exploratory resting-state analyses, whether any language-associated clade (a group of components with similar time courses) changed its coupling with sensory cortex under psilocybin, and whether participants with higher EDI scores showed larger connectivity changes.

**Figure 1.**
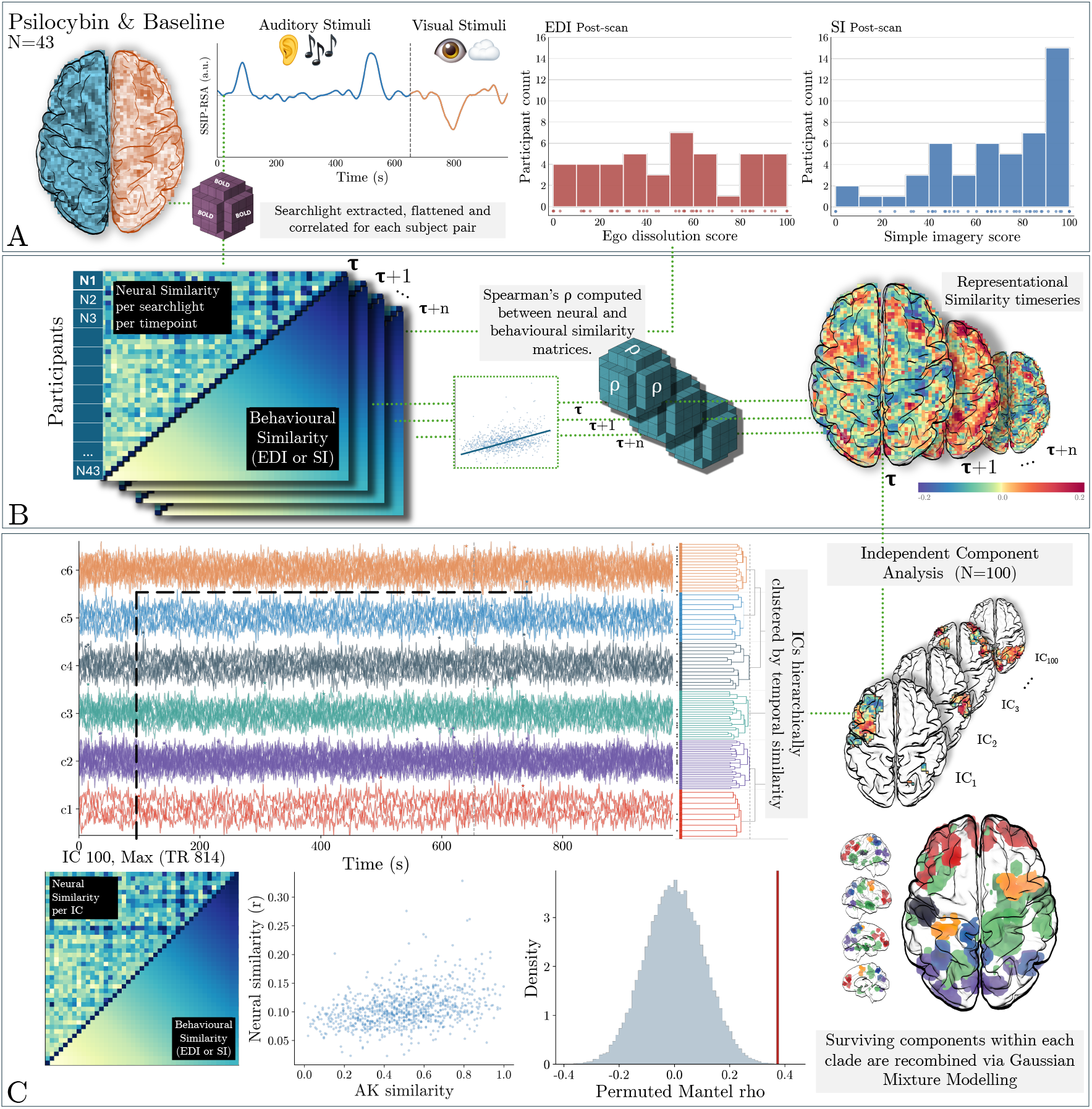
Time-resolved Spatial Inter-Subject Representational Similarity Analysis (SIS-RSA) pipeline. (A) Hyperaligned multivoxel patterns were extracted for each participant from overlapping searchlights at every TR across the concatenated music-listening and cloud-gazing tasks. Histograms (right) show the distributions of EDI and elementary imagery (SI) scores. (B) At each searchlight and TR, pairwise Pearson correlations between participant patterns formed a neural-similarity matrix. Spearman correlation of its upper triangle with the directional rank-sum behavioural matrix yielded one SIS-RSA coefficient; repetition over space and time produced a four-dimensional SIS-RSA image. (C) We decomposed each image with MELODIC into 100 spatial independent components (ICs) and their time courses. For each IC, the best positive-loading mask at the TR of peak expression was tested against a Mantel null distribution from 50,000 participant-label permutations. All 100 IC time courses were grouped into clades by Ward clustering, and the positive Gaussian-mixture-model (GMM) cores of ICs surviving q<.01 were combined within each clade.

## Methods

### Participants, parent study and phenomenological measures

Data were drawn from the 62-person PsiConnect cohort (ACTRN12621001375842), a multimodal study of context-dependent brain and behavioural dynamics before and after psilocybin administration (Novelli et al., 2026; Stoliker et al., 2026). Healthy adults aged 18 to 55 years received a single 19 mg oral dose of psilocybin. Recruitment, clinical screening, drug administration, monitoring and the full acquisition schedule are described in the parent report (Stoliker et al., 2026) and data descriptor (Novelli et al., 2026). The study was approved by the Monash University Human Research Ethics Committee (MUHREC 27181), and all participants provided written informed consent. The parent design included immediate-entry and meditation-training arms; participants were pooled in the present analysis because the dose, imaging sessions, naturalistic runs and post-dose measures used here were common across arms.

The present analysis used the first post-scan administration of the eight-item Ego Dissolution Inventory (EDI; Nour et al., 2016), completed immediately after scanning (approximately 120 minutes post-dose), and the elementary imagery dimension of the 11-Dimensional Altered States of Consciousness questionnaire (11D-ASC; Studerus et al., 2010), completed at the end of the day after participants had returned to baseline. Elementary (simple) Imagery, abbreviated SI throughout the manuscript, measures recalled elementary visual phenomena such as geometric patterns and colours. We selected it as a phenomenological comparator because elementary imagery is a prominent but conceptually distinct aspect of the psychedelic state. We did not assume that SI was process-pure or statistically independent of EDI, and no inference was based solely on significance for one measure and non-significance for the other.

Participants contributing to each measure-specific analysis had complete, common stimulus-aligned music and cloud-viewing series and a directly observed score for that measure. This yielded 43 participants for EDI and 49 for SI; 42 contributed to both analyses, and their union comprised the 50-participant hyperalignment pool (age 19–53 years; Table 1).

**Table 1.**
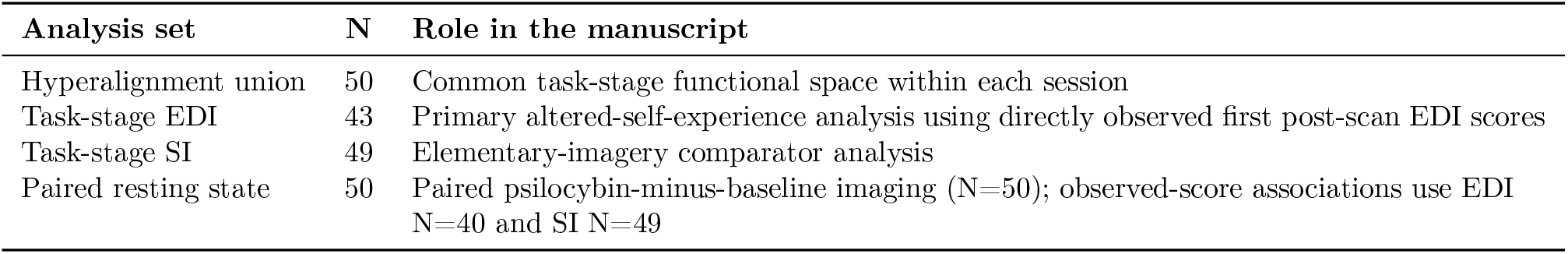
Samples used for hyperalignment, task SIS-RSA and resting-state connectivity. The union of the observed-score EDI and SI task samples defined the session-specific hyperalignment spaces. Resting-state imaging inference used all 50 paired scans passing quality control; behavioural correlations used the subsets with observed EDI or SI scores.

| Analysis set | N | Role in the manuscript |
| --- | --- | --- |
| Hyperalignment union | 50 | Common task-stage functional space within each session |
| Task-stage EDI | 43 | Primary altered-self-experience analysis using directly observed first post-scan EDI scores |
| Task-stage SI | 49 | Elementary-imagery comparator analysis |
| Paired resting state | 50 | Paired psilocybin-minus-baseline imaging (N=50); observed-score associations use EDI N=40 and SI N=49 |

### MRI acquisition, naturalistic stimuli and released preprocessing

The two naturalistic acquisitions supplied a common time-varying sensory sequence against which participant-specific neural representations could be compared. Images were acquired on a Siemens Skyra 3 T scanner with a 32-channel receive head coil. Functional data used a four-echo echo-planar sequence with repetition time (TR) .91 s, echo times 12.60, 29.23, 45.86 and 62.49 ms, 3.2-mm isotropic voxels and a 64 × 64 acquisition matrix. Participants completed the same runs at baseline, approximately one week before dosing, and during the psilocybin session, approximately 80 minutes after dosing. The music run contained two uninterrupted instrumental tracks: Spacecraft and Lauge, “Sunset from the Ethereal Shore (Lauge Rework)”, followed by Lauge and Applefish, “Sollys (Applefish Rework)”. The visual run presented silent, slowly moving cloud imagery. The common stimulus-aligned analysis windows contained 728 music volumes and 372 cloud-viewing volumes. Removing the first ten volumes of every run left 718 music and 362 cloud-viewing volumes.

Anatomical and functional preprocessing used the PsiConnect pipeline (Novelli et al., 2026). fMRIPrep 22.0.2 corrected and spatially normalised the functional data to the MNI152NLin2009cAsym common anatomical space (Esteban et al., 2019). For the multiecho acquisitions, tedana 0.0.12 optimally combined the echoes and used independent-component analysis to identify components classified as non-BOLD (DuPre et al., 2021). The released non-global-signal-regressed BOLD series then underwent a nuisance general linear model in SPM12 containing framewise displacement, mean white-matter and cerebrospinal-fluid signals, the time courses of tedana-classified non-BOLD components and an intercept. The present analysis used these released nuisance-cleaned images and did not repeat preprocessing.

### Analysis overview

The primary contrast related EDI scores to psilocybin-session neural data. We specified the primary prediction from pilot analyses of an independent LSD dataset before analysing the psilocybin data, but did not preregister it. We used three identically processed control contrasts to test alternative explanations: baseline–EDI to ask whether the ordering associated with later EDI was already present before dosing, psilocybin–SI to ask whether a distinct perceptual dimension of the psychedelic state produced the same association, and baseline–SI as the baseline counterpart of psilocybin–SI. We refer to these analyses as psilocybin–EDI, baseline–EDI, psilocybin–SI and baseline–SI. Each comprised hyperalignment, time-resolved SIS-RSA, independent-component decomposition, component-wise Mantel testing, temporal clustering and meta-analytic decoding (Figure 1). A separate resting-state acquisition was then used to test connectivity changes in the task-defined psilocybin– EDI clades.

### Concatenated searchlight hyperalignment

We hyperaligned the data because functionally corresponding multivoxel patterns occupy idiosyncratic anatomical configurations across people. For each session separately, music and cloud-viewing data were concatenated to form a 1,080-volume series for each participant and z-standardised across the complete concatenated run. Searchlight hyperalignment, implemented in PyMVPA2, used a three-voxel radius and three-voxel sparse spacing to learn a high-dimensional common information space from the 50-participant union (Table 1) without access to EDI or SI labels (Haxby et al., 2011, 2020; Guntupalli et al., 2016). The transformations were applied to each participant and the aligned series were divided back into music and cloud portions. The non-invariant common masks contained 32,411 voxels under psilocybin and 32,392 voxels at baseline. We learned a separate common space for each session.

### Spatial inter-subject representational similarity analysis

We used SIS-RSA because EDI-related neural convergence could be expressed by different systems at different moments of a naturalistic sequence (Ma and Skipper, 2025). At each TR and searchlight, every participant contributed a vector containing the local voxel values. Pearson correlation between every participant pair produced an N × N neural-similarity matrix. Spearman correlation between the unique off-diagonal entries of that matrix and the corresponding entries of the directional behavioural matrix yielded one coefficient, assigned to the searchlight centre. Repeating this calculation over the brain and the concatenated music and cloud-viewing series produced a four-dimensional SIS-RSA coefficient image.

SIS-RSA used a fixed spherical neighbourhood containing voxel centres within 2.5 voxels (approximately 8.0 mm) of each searchlight centre, with a mean of 59.3 voxels inside the common mask. This geometry was applied identically to all four analyses and was distinct from the three-voxel neighbourhood used for hyperalignment.

The directional behavioural matrix was constructed from ranks of the observed EDI or SI scores. For N participants with ranks r_i and r_j, the off-diagonal entry was B_ij=(r_i+r_j)/(2N). The off-diagonal entries were then linearly rescaled to the unit interval. The diagonal was set to one, and only the strict upper triangle, excluding the diagonal, entered SIS-RSA and Mantel tests. The model therefore orders pairs by joint score rank: high–high pairs receive the largest values, mixed pairs intermediate values and low–low pairs the smallest. This ordering implements the Anna Karenina model of Finn et al. (2020), which we preferred to their nearest-neighbour model because the latter predicts similarity from the distance between two participants’ scores and would therefore treat any two low scorers with adjacent scores as convergent. For EDI, the directional hypothesis was that self-related representations are normally idiosyncratic, so participants reporting greater dissolution would show greater interparticipant neural convergence within territories supporting selfhood. A positive SIS-RSA coefficient indicates increasing neural similarity along this joint-score ordering.

### Independent-component decomposition and component-wise testing

Because the time-resolved SIS-RSA images could contain different behaviour-related systems at different moments, we used FSL MELODIC to recover recurring spatial modes, decomposing each four-dimensional image into 100 spatial independent components (ICs) and associated time courses (Beckmann and Smith, 2004). The dimensionality matched the high-dimensional naturalistic SIS-RSA implementation of Ma and Skipper (2025), was held constant across all four analyses and was not selected from the decoding results. Each IC was evaluated at the TR at which the absolute value of its IC time course was maximal. Its spatial map was divided by its maximum absolute loading, and nine positive masks were generated at thresholds from .1 to .9. Masks with fewer than 50 voxels in total were omitted so that pair-wise spatial-pattern correlations were not estimated from very small supports. At that peak-expression TR, the eligible spatial mask producing the greatest positive Spearman association with the directional behavioural matrix was retained. Only spatial support was optimised against the behavioural matrix; the TR was fixed by the IC time course alone.

We then used a one-tailed Mantel test with 50,000 participant-label permutations to test whether each selected component mask was more positively related to the directional behavioural matrix than expected after breaking participant correspondence. Benjamini–Hochberg correction was applied across the 100 ICs separately within each analysis (Benjamini and Hochberg, 1995), with q<.01 defining the retained set. The spatial support was selected once from the observed data and was not reselected within every permutation; p values are therefore conditional on that selected support and are treated as discovery-oriented component statistics.

### Clade construction and GMM-core maps

We grouped the 100 ICs into coalitions whose SIS-RSA expression rose and fell at similar moments, so that their anatomy could be merged and decoded. Ward hierarchical clustering was applied to correlation distances between all 100 IC time courses. The dendrogram was cut at the 95th percentile of merge heights as a pragmatic balance between very broad coalitions and an unwieldy number of narrow groups, and the same rule was applied to all four analyses; it yielded six clades in each. Only q<.01 ICs assigned to a clade contributed to its spatial map. For each contributing IC separately, a two-component Gaussian mixture model (GMM) was fitted to the complete voxel-loading distribution. The lower-mean component represented background or low loadings, while the higher-mean component identified the positive-loading core without another fixed loading threshold. Positive voxels with posterior probability greater than .5 for the higher-mean component were retained, following the GMM procedure of Ma and Skipper (2025). This step defined spatial support; the behavioural association was tested only at the component level. At each voxel, retained original loadings were averaged across the significant ICs whose cores included that voxel. Each clade was thresholded at its 75th positive-loading percentile for display only (Figures 2 and 3).

**Figure 2.**
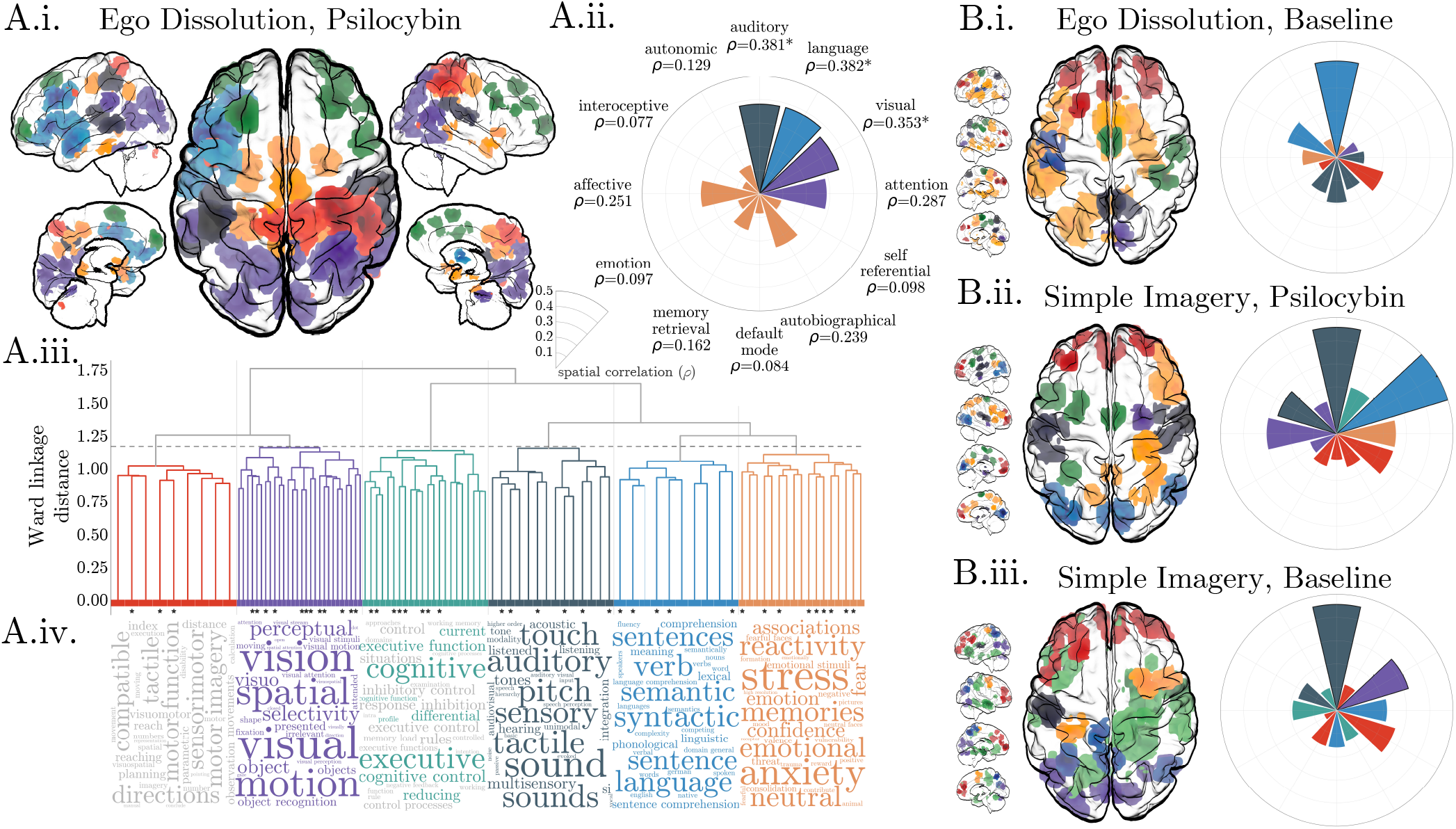
Spatial organisation and functional decoding of the six temporal clades. (A) Psilocybin–EDI: (i) merged clade maps displayed above the within-map 75th loading percentile, (ii) the largest positive clade correlation for each displayed a priori Neurosynth term, coloured by the winning clade, (iii) the Ward dendrogram with the fixed cut and significant ICs marked by asterisks, and (iv) complete-vocabulary cognitive terms for each clade, aligned with its dendrogram branch (c1–c6, left to right), with non-significant terms desaturated. (B.i–iii) Merged clade maps and a priori term correlations, as in A.i–ii, for baseline–EDI (B.i), psilocybin–SI (B.ii) and baseline–SI (B.iii); terms occupy the same polar positions as in A.ii. At each voxel, clade-map intensity is the mean MELODIC spatial z score across the significant ICs whose positive-GMM cores include that voxel. Each clade is independently scaled from the 75th percentile of its non-zero values to its within-clade maximum, so colour intensity is comparable only within a clade. Thresholding was for display only; decoding used the complete continuous maps. Exact display limits are reported in Supplementary Data S1.

**Figure 3.**
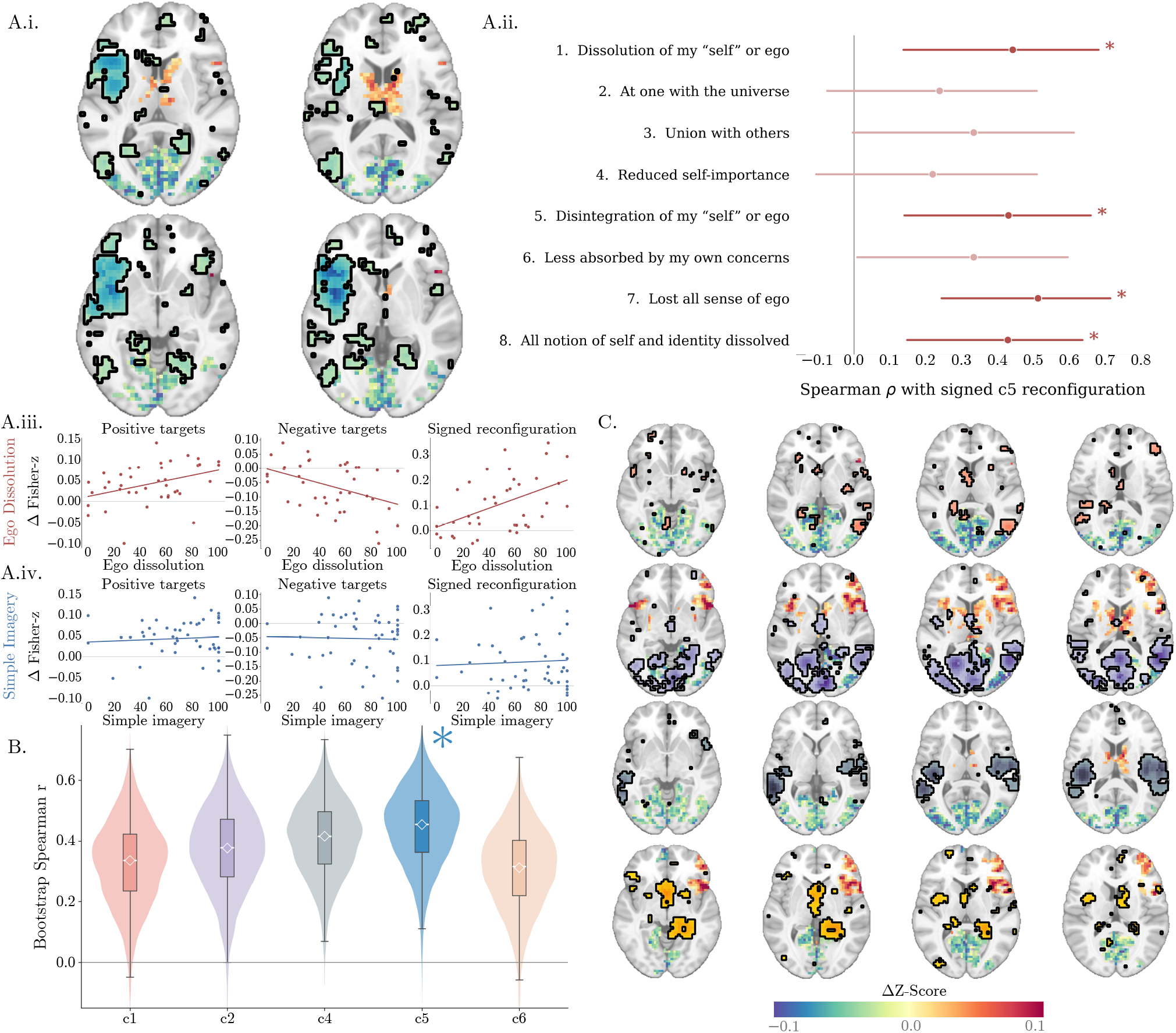
Psilocybin-related resting-state connectivity changes in task-defined ego-dissolution clades. (A) Language-associated c5: (i) seed and FWER-corrected group-mean connectivity change; (ii) correlations between the positive-minus-negative score and individual EDI items (N=40; bars are 95% bootstrap intervals; ^*^Holm-adjusted p<.05); and mean changes in increasing targets, decreasing targets and their difference plotted against (iii) EDI and (iv) SI. Lines are descriptive least-squares fits. (B) Bootstrap distributions of the EDI correlation with the positive-minus-negative score across clades; diamonds show observed Spearman ρ and the asterisk marks c5, the only association surviving Bonferroni correction across 15 tests. (C) Seeds and corrected connectivity changes for c1, c2, c4 and c6; c3 had no corrected targets. Axial sections in A.i and C are at MNI z=™5.0, 1.5, 8.0 and 14.5 mm. Maps show the group-mean psilocybin-minus-baseline change in Fisher-transformed correlation (Δz), from ™.10 to .10. Seeds are thresholded at their 75th percentile for display only.

Anatomical summaries were computed descriptively from each complete positive-GMM clade, rather than the 75th-percentile display masks. Each map was intersected on its exact analysis grid with the Schaefer 1,000-parcel, seven-network cortical atlas (Schaefer et al., 2018) and the AFNI-distributed FreeSurfer MNI152NLin2009cAsym subcortical segmentation (Fischl et al., 2002); the subcortical atlas was resampled to the clade grid using nearest-neighbour interpolation. For each analysis and clade, we report voxel counts and percentages of clade support and the mean and maximum MELODIC spatial z within each cortical parcel/network and subcortical label. These summaries are reported in Supplementary Table S1 and Supplementary Data S1.

### Meta-analytic decoding and spatial nulls

Meta-analytic decoding supplied provisional functional labels for the clade maps. Adapting the narrative- and embodied-self terms used by Ma and Skipper (2025), we selected language, speech perception, speech production, autobiographical, memory retrieval, affective, autonomic, emotion, interoceptive and self-referential maps. Auditory and visual were added as stimulus and sensory comparators, attention as a task-engagement comparator and default mode as the principal competing-network account, giving 14 Neurosynth version 7 association maps in total (Yarkoni et al., 2011). The exploratory complete-vocabulary analysis tested 3,228 terms represented in at least 50 studies.

Clade and Neurosynth maps are spatially auto-correlated, so treating their voxels as independent would underestimate null variation. Pearson correlation quantified the spatial correspondence between each continuous clade map and each Neurosynth map. For every clade, 10,000 BrainSMASH surrogate maps preserved the empirical variogram—the relationship between map similarity and spatial distance—on the native 3.2-mm grid while disrupting anatomical correspondence, using 1,500 nearest neighbours (Burt et al., 2020). Two-sided empirical p values compared the absolute observed clade–term correlation with the surrogate distribution. Benjamini–Hochberg correction covered all 84 a priori clade-by-term tests separately within each analysis; q<.01 was the primary threshold. Complete-vocabulary correction was performed separately within each clade across 3,228 terms, with q<.05 treated as exploratory. For Figure 2 only, each clade map was displayed above its within-map 75th positive-loading percentile for legibility; every decoding test used the complete continuous GMM-core map.

### Resting-state sample

To test whether task-defined clades changed their functional connectivity under psilocybin, and whether participants reporting greater ego dissolution showed larger changes, we analysed a separate pair of baseline and psilocybin resting-state scans without hyperalignment, using seeds defined from the task data alone. We used the six psilocybin–EDI clades as seeds, with c5, whose map corresponded to the Neurosynth language map, as the primary seed and the other five as comparators, and related the same connectivity measures to both EDI and SI. Standard scans contained 504 volumes; the first five were identically zero and discarded, leaving 499 candidate volumes before connectivity-specific censoring.

The resting-state series had already undergone the released PsiConnect preprocessing described above. For connectivity estimation, we additionally censored high-motion or unstable frames: framewise displacement greater than .30 mm together with one preceding and two following frames; standardised DVARS greater than 1.5 and existing fMRIPrep motion-outlier indicators at the flagged frame only; and non-steady-state indicators at the flagged frame only. Each session had to retain at least 330 volumes— approximately five minutes—and have no more than 34% of the frames after the discarded initial volumes censored; both sessions had to pass. Fifty participants met these paired imaging criteria. Behavioural correlations used every participant in this imaging cohort with an observed score: EDI N=40 and SI N=49.

### Task-derived seeds and voxelwise resting-state connectivity

We converted each clade’s full positive GMM core, within the common grey-matter mask, into a binary seed, so that every positive-loading voxel contributed equally to the seed time course; the 75th-percentile seeds shown in the figures are for display only. For each participant and session, we averaged the BOLD signal across seed voxels over the retained frames, correlated this time course with every grey-matter voxel outside the seed (Pearson) and converted the correlations to Fisher z.

We tested each participant’s psilocybin-minus-baseline difference in Fisher-z connectivity against zero with AFNI 3dttest++. Voxelwise multiplicity was controlled with Equitable Thresholding and Clustering (ETAC; Cox, 2017), using two-sided inference, nearest-neighbour setting 2, 0-, 4- and 8-mm blur levels, and voxelwise thresholds of .01, .005, .002 and .001 within the exact grey-matter-minus-seed target mask. Every clade used the identical within-map family-wise error rate (FWER) of 5%.

The corrected map was separated into regions whose seed connectivity increased under psilocybin and regions whose seed connectivity decreased, according to the sign of the group t statistic. For each participant and clade, we averaged the psilocybinminus-baseline Fisher-z connectivity change over the increasing regions and separately over the decreasing regions. To express each participant’s change across the whole corrected pattern as one number, with increases and decreases weighted equally, we subtracted the decreasing-region mean from the increasing-region mean to give a positive-minus-negative connectivity score. These three participant-level measures were Spearman-correlated with observed EDI and SI. C3 had no corrected voxels, so we applied Bonferroni correction across the remaining 15 EDI tests (five clades by three measures) to judge the c5 association within the full family of clade tests.

## Results

### Stimulus-driven synchrony remained under psilocybin

We used conventional intersubject-correlation (ISC) analyses (Supplementary Figure 1) to check two assumptions of the main analysis. First, because we aligned and compared participants at the same moment of the same stimulus, their responses had to stay partly shared; mean whole-brain ISC stayed positive under psilocybin, so a stimulus-driven component survived (Supplementary Figure 1A–B). Second, if psilocybin made participants track the stimuli more closely, synchrony should have risen in both tasks, and periods of shared response should have lasted longer. Instead, ISC fell during music and rose during cloud viewing, and during music, periods of local synchrony were shorter (the autocorrelation of local synchrony first crossed zero after 5.56 s on average, against 7.14 s at baseline; Supplementary Figure 1C).

### SIS-RSA identified six clusters of IC time courses in each analysis

In each analysis, we decomposed the time-resolved SIS-RSA images into 100 independent components (ICs) and tested each IC by permuting participant labels 50,000 times (Figure 1C). Observed EDI and SI scores were available for 43 and 49 participants, respectively. Forty-two participants provided both measures, which were weakly correlated (Spearman ρ=.101, p=.523). After Benjamini– Hochberg correction across 100 ICs within each analysis (q<.01), 43 ICs were retained for psilocybin–EDI, 31 for psilocybin–SI, 29 for baseline–EDI and 32 for baseline–SI. All retained component-level associations were positive.

Ward clustering of the 100 IC time courses yielded six clades in each analysis. Here, a clade denotes a cluster of ICs with similar time courses. Only ICs meeting q<.01 in the component-level Mantel tests contributed to its spatial map. In psilocybin–EDI, c1 spanned right-dominant sensorimotor cortex; c2, occipital cortex; c3, right-dominant lateral prefrontal cortex; c4, superior-temporal/opercular and somatomotor cortex; c5, a distributed left-lateralised fronto-temporal map; and c6, medial-temporal, limbic and posterior-medial regions. In the meta-analytic decoding reported below (Figure 2A; Supplementary Table S1), these maps corresponded most closely with visual (c2), executive (c3), auditory (c4), language (c5), and affective and mnemonic (c6) terms, which we use as the clades’ labels. C5 comprised five retained ICs (mean component-level Mantel ρ=.296), and its complete positive GMM-core map contained 5,701 voxels. It peaked in dorsal pars opercularis of the left inferior frontal gyrus, near the inferior frontal junction (MNI [™45, 16, 30]), and extended across adjacent lateral prefrontal cortex, the frontal operculum and insula, and left lateral temporal cortex and temporal pole. Supplementary Table S1 and Supplementary Data S1 report anatomical summaries for all 24 analysis-by-clade maps.

### Comparator analyses contained distinct anatomical clade structures

We ran the three comparator analyses to test alternative explanations of the primary result: whether pairs reporting more ego dissolution already had more similar activity before dosing (baseline–EDI), and whether elementary imagery, a perceptual dimension of the psychedelic state, showed the same association (psilocybin–SI, with baseline–SI as its baseline counterpart). We describe their anatomy here and their decoding in the next subsection. The three comparator analyses also produced anatomically differentiated clades (Figure 2B.i–iii; Supplementary Table S1). In psilocybin–SI, c1 spanned bilateral prefrontal cortex; c2 was a compact and strongly right-lateralised limbic–subcortical map encompassing amygdala, hippocampus, orbitotemporal cortex and putamen; c3 was strongly left-lateralised across premotor and frontoparietal cortex; c4 occupied temporal–opercular and somatomotor cortex; c5 was centred on bilateral occipital cortex; and c6 was a right-lateralised frontoparietal map. These clades differed considerably in scale: c2 contained one significant IC and 489 voxels, whereas c6 contained eight ICs and 4,649 voxels.

Baseline–EDI comprised frontal c1, occipital c2, sensorimotor c3, posterior-medial c4 including precuneus and posterior cingulate cortex, a strongly left-lateralised temporal–opercular c5, and a large distributed cortical c6 together with hippocampal, thalamic, striatal and cerebellar structures. Baseline–SI similarly contained frontal c1, occipital c2, a large distributed c3, a strongly left-lateralised temporal–opercular c4, posterior parietal c5 and right-dominant premotor c6. The largest comparator maps were baseline–EDI c6 (15 significant ICs; 13,364 voxels) and baseline–SI c3 (seven ICs; 11,892 voxels). Exact support sizes, peaks, cortical-network composition, parcel overlaps and subcortical overlaps are reported in Supplementary Table S1 and Supplementary Data S1.

### Psilocybin–EDI c5 showed the strongest language-map correspondence

To test the primary prediction, we correlated every clade map with the Neurosynth language map and assessed each correlation against 10,000 surrogate maps with the same spatial autocorrelation as the clade map. If participants reporting more ego dissolution converge in language-associated cortex under psilocybin, a psilocybin–EDI clade should match the language map more closely than the best-matching clade in any comparator analysis. The psilocybin– EDI c5 map correlated with the Neurosynth language meta-analytic association map (Pearson r=.382, two-sided spatial-null p<.001, q=.003 after correction across 84 clade-by-term tests; Table 2; Figure 2A.ii). Among the 84 clade-by-term correlations in psilocybin–EDI, only auditory c4 matched this value (r=.381); the a priori terms matching c6, c1 and c3 most closely were affective (r=.251), attention (r=.145) and memory retrieval (r=.088; Supplementary Data S2). The maximum positive language-map correlations in the comparator analyses were numerically smaller: psilocybin–SI c3, r=.208, q=.028; baseline–EDI c6, r=.055, q=.794; and baseline–SI c1, r=.117, q=.343. Only psilocybin–EDI met the primary q<.01 threshold (Figure 2B.i–iii).

**Table 2.** Largest positive clade correspondence for each displayed a priori Neurosynth term in each analysis. Entries report the winning clade, Pearson r and Benjamini–Hochberg-adjusted spatial-null q across all 84 clade-by-term tests within that analysis. Asterisks denote q<.01. Supplementary Data S2 reports all 336 a priori analysis-by-clade-by-term tests, including the two speech terms omitted from this compact display.

| A priori term | Psilocybin–EDI | Baseline–EDI | Psilocybin–SI | Baseline–SI |
| --- | --- | --- | --- | --- |
| Language | **c5; $r=.382$; $q=.003^*$** | c6; $r=.055$; $q=.794$ | c3; $r=.208$; $q=.028$ | c1; $r=.117$; $q=.343$ |
| Auditory | **c4; $r=.381$; $q=.003^*$** | **c5; $r=.415$; $q=.008^*$** | **c4; $r=.458$; $q=.002^*$** | **c4; $r=.456$; $q=.004^*$** |
| Visual | **c2; $r=.353$; $q=.003^*$** | c2; $r=.096$; $q=.641$ | **c5; $r=.540$; $q=.002^*$** | **c2; $r=.322$; $q=.004^*$** |
| Attention | c2; $r=.287$; $q=.012$ | c4; $r=.119$; $q=.627$ | c6; $r=.255$; $q=.025$ | c5; $r=.217$; $q=.109$ |
| Autobiographical | c6; $r=.239$; $q=.030$ | c4; $r=.146$; $q=.466$ | c1; $r=.150$; $q=.277$ | c3; $r=.138$; $q=.333$ |
| Memory retrieval | c6; $r=.162$; $q=.179$ | c4; $r=.161$; $q=.283$ | c1; $r=.146$; $q=.215$ | c1; $r=.160$; $q=.173$ |
| Self-referential | c6; $r=.098$; $q=.559$ | c1; $r=.211$; $q=.112$ | c1; $r=.254$; $q=.012$ | c1; $r=.264$; $q=.015$ |
| Default mode | c6; $r=.084$; $q=.559$ | c4; $r=.195$; $q=.022$ | c1; $r=.113$; $q=.294$ | c5; $r=.159$; $q=.109$ |
| Affective | c6; $r=.251$; $q=.030$ | c6; $r=.148$; $q=.466$ | c2; $r=.302$; $q=.025$ | c3; $r=.191$; $q=.173$ |
| Emotion | c6; $r=.097$; $q=.421$ | c1; $r=.081$; $q=.627$ | c2; $r=.120$; $q=.277$ | c1; $r=.057$; $q=.679$ |
| Autonomic | c6; $r=.129$; $q=.309$ | c6; $r=.085$; $q=.641$ | c2; $r=.145$; $q=.292$ | c3; $r=.101$; $q=.448$ |
| Interoceptive | c6; $r=.077$; $q=.559$ | c5; $r=.222$; $q=.013$ | c4; $r=.265$; $q=.002^*$ | c4; $r=.174$; $q=.070$ |

### A priori sensory correspondence provided positive controls

Because every participant heard the same music and saw the same clouds, we expected some clade in every analysis to match the auditory and visual maps, and we used these matches to check our decoding. Auditory-map correlations met q<.01 in all four analyses (r=.381–.458). Within psilocybin–EDI, auditory and speech-perception correspondence occurred in c4 (r=.381, q=.003 and r=.310, q=.008, respectively), whereas language correspondence occurred in c5. Visual-map correspondence met q<.01 in psilocybin–EDI c2 (r=.353, q=.003), psilocybin–SI c5 (r=.540, q=.002) and baseline–SI c2 (r=.322, q=.004), but not baseline–EDI. Neither the self-referential nor default-mode map met q<.01 in any analysis (Table 2; Supplementary Data S2).

In the exploratory complete-vocabulary Neurosynth analysis of the psilocybin–EDI clades (Figure 2A.iv), the highest-ranked corrected terms included visual, vision and motion for c2; executive and cognitive control for c3; sound, auditory, tactile and pitch for c4; syntactic, verb, language, sentences and semantic for c5; and stress, anxiety, reactivity, memories and emotional for c6. No term met q<.05 for c1, a predominantly sensorimotor clade whose nearest neighbour in the dendrogram was visual c2 (Figure 2A.iii).

Psilocybin–SI showed a different organisation. Corrected complete-vocabulary terms associated c1 with prospective cognition and choice, c2 with fear and face-processing paradigms, c3 with planning and motor imagery, c4 with pitch and auditory processing, c5 with visual motion and c6 with task difficulty. Baseline decoding was sparser. In baseline–EDI, complete-vocabulary terms survived q<.05 only for sensorimotor c3 and auditory c5; in baseline–SI, they survived for executive/rule-related c1, visual and object-recognition c2, and auditory c4. No complete-vocabulary term survived correction in the remaining baseline clades. Supplementary Table S1 provides a concise anatomy-and-decoding summary; Supplementary Data S1 and S3 provide the complete anatomical overlaps and term-level statistics.

### Resting-state connectivity of psilocybin–EDI c5 differed between baseline and psilocybin

Because language shapes perception, we next asked whether psilocybin changed how the language-associated clade couples with sensory cortex, using c5 as the primary seed in separate resting-state scans. In the paired resting-state imaging cohort (N=50), all six psilocybin–EDI clades were tested using identical two-sided ETAC procedures with a within-map family-wise error rate of .05. Significant connectivity changes were detected for c1, c2, c4, c5 and c6; none was detected for c3 (Table 3; Figure 3C). For c5, 2,161 voxels survived correction: connectivity increased in 317 and decreased in 1,844. The largest contiguous region contained 1,847 voxels, of which 1,838 showed decreased connectivity, and peaked in medial occipital cortex (MNI [0, ™81, 14]; group-mean psilocybin-minus-baseline change in Fisher z at the peak, Δz=™.115). A 271-voxel region, of which 265 voxels showed increased connectivity, peaked in right thalamus ([10, ™20, 11]; peak Δz=.062); smaller corrected regions were present in medial and right frontal cortex (Figure 3A.i).

**Table 3.** Matched resting-state connectivity results for all six psilocybin–EDI clades. Corrected targets come from identical two-sided within-map FWER 5% ETAC tests. Behavioural columns report the correlation with each participant’s increasing-target mean minus decreasing-target mean; Bonferroni p values use all 15 estimable EDI clade-by-measure tests.

| Psilocybin–EDI clade | Corrected target voxels, n (increase/decrease) | EDI $\rho$ | Raw p | Bonferroni p (15 tests) | SI $\rho$ | Raw p |
| --- | --- | --- | --- | --- | --- | --- |
| c1 sensorimotor | 3,102 (46/3,056) | .337 | .0337 | .506 | −.029 | .844 |
| c2 visual | 3,669 (2,487/1,182) | .377 | .0164 | .246 | .140 | .338 |
| c3 lateral prefrontal | 0 | — | — | — | — | — |
| c4 auditory | 2,577 (197/2,380) | .416 | .00756 | .113 | −.001 | .997 |
| c5 distributed frontotemporal / language | 2,161 (317/1,844) | .455 | .00321 | .0482 | .008 | .957 |
| c6 medial temporal | 2,750 (1,279/1,471) | .313 | .0493 | .740 | .227 | .117 |

### C5 connectivity change was associated with EDI but not SI

If the change in c5’s connectivity reflects the disruption that accompanies ego dissolution, participants reporting more ego dissolution should show larger changes, and elementary imagery should predict those changes less well. We then correlated each participant’s connectivity change with EDI and SI in exploratory analyses, using the increasing-target mean, the decreasing-target mean and the positive-minus-negative score, which expresses change across the whole corrected pattern.

Among participants with observed EDI scores (N=40), mean change in c5 increasing targets correlated positively with EDI (ρ=.380, p=.0156), whereas mean change in decreasing targets correlated negatively (ρ=™.338, p=.0328; Figure 3A.iii). The positive-minus-negative score showed the largest association (ρ=.455, p=.00321) and was the only one to survive Bonferroni correction across the 15 estimable clade-by-measure EDI tests (adjusted p=.0482). Positive-minus-negative scores for the other four estimable clades were also positively associated with EDI before correction (ρ=.313–.416, raw p=.0076–.049), most strongly for c4 (ρ=.416) and c2 (ρ=.377), but none survived Bonferroni correction (Table 3; Figure 3B). For c4 and c2, EDI correlated with mean change in increasing targets at ρ=.390 and ρ=.256 and with mean change in decreasing targets at ρ=™.300 and ρ=™.390, respectively (raw p=.013–.110). The c5 score was not associated with SI (N=49; ρ=.008, p=.957; Figure 3A.iv), and neither were those of the other clades (all p≥.117; Table 3).

In an exploratory analysis, we related the c5 positive-minus-negative score to the eight individual EDI items in the same N=40 cohort (Figure 3A.ii). Four associations survived Holm correction: dissolution of the ‘self’ or ego (ρ=.442, adjusted p=.030), disintegration of the ‘self’ or ego (ρ=.430, adjusted p=.034), loss of all sense of ego (ρ=.512, adjusted p=.006), and dissolution of all notions of self and identity (ρ=.428, adjusted p=.034). Associations with unity, reduced self-importance and reduced absorption in personal concerns did not survive correction (ρ=.219–.334, adjusted p=.142–.277).

## Discussion

### A language-associated correlate of ego dissolution

Under psilocybin, participants with higher EDI scores had more similar fMRI activity patterns to one another in a distributed, left-lateralised network (c5), which matched the Neurosynth language map (r=.382) more closely than the best-matching clade in any comparator analysis (r=.055–.208; Table 2; Figure 2). At rest, this network’s connectivity decreased with posterior visual cortex and increased with thalamic and frontal targets (Figure 3A). A connectivity score contrasting these increases with the decreases was higher in participants reporting more ego dissolution (ρ=.455) and unrelated to elementary imagery (ρ=.008), so among the experiences we measured, the association was specific to loss of self.

Although the language-associated network (c5) extended beyond classical language regions into parcels assigned to default, control and salience/ventral-attention networks (Supplementary Data S1), its regions have established roles in language or semantic processing: lateral temporal cortex lies where semantic and default-network systems meet (Binder et al., 2009; Seghier et al., 2010); the temporal pole is a proposed amodal semantic hub (Patterson et al., 2007); and the striatum, thalamus and cerebellum contribute to lexical selection and semantic access (Kotz and Schwartze, 2010; Crosson, 2013). Left inferior frontal involvement could instead reflect domain-general cognitive control, but a separate, right-lateralised prefrontal clade decoded to executive and cognitive-control terms, and the Neurosynth attention map correlated best with the visual clade (Table 2; Supplementary Table S1).

Medial-temporal, parahippocampal–retrosplenial and posterior cingulate regions previously associated with ego dissolution (Muthukumaraswamy et al., 2013; Lebedev et al., 2015; Carhart-Harris et al., 2016), and the amygdala, whose coupling psilocybin alters (Stoliker et al., 2024), fell instead in a separate affective and mnemonic clade (c6). This separation is consistent with proposals that ego dissolution comprises partly dissociable changes in bodily self-consciousness, agency, first-person perspective and narrative selfhood (Millière, 2020; Stoliker et al., 2022; Sleight et al., 2023). To network-level accounts of ego dissolution, which invoke increased global connectivity (Tagliazucchi et al., 2016), altered coupling between anticorrelated systems (Stoliker et al., 2023) and stronger prefrontal–thalamic connectivity (Coleman et al., 2025), we add the disruption of a cognitive process: the incorporation of person-specific semantic and autobiographical context into ongoing experience.

### Interpreting interparticipant convergence

Psilocybin has previously made whole-brain functional connectivity more similar between participants while leaving each participant identifiable (Tolle et al., 2024), and the same convergence occurred during ritual ayahuasca use, where a shared religious community could explain the effect (Mallaroni et al., 2024). Our participants were scanned individually, outside any ritual context, so a shared setting cannot explain their convergence, which was larger in pairs reporting more ego dissolution and centred on language-associated cortex. In the same cohort, psilocybin also aligned brain dynamics with the experimental context, more strongly in participants reporting deeper self- and boundary-dissolving experiences (Stoliker et al., 2026). Both results are consistent with the account proposed here: if personal associations shape responses less, responses follow the shared stimulus more closely, so they align more with the context and resemble one another more.

Alternatively, because intersubject correlation increases with attention and engagement (Ki et al., 2016), highly absorbed participants may simply have tracked the music more closely. Under psilocybin, however, mean whole-brain ISC decreased during music, when participants also fell out of local synchrony sooner (most strongly outside c5), and increased during cloud viewing (Supplementary Figure 1A–C), so a global increase in stimulus tracking is unlikely to explain the convergence. Inattention, conversely, weakens shared stimulus-driven responses (Ki et al., 2016), so if high scorers had attended less, they should have converged less in sensory cortex, yet high-scoring pairs also converged in the auditory (c4) and visual (c2) clades.

Convergence could also reflect how participants interpret stimuli. What a word or event denotes is shared across people, while the connotations it evokes remain personal. However, instrumental music and drifting clouds have few denotations. Because psychedelics heighten the perceived meaningfulness of ordinary experience (Hartogsohn, 2018), participants under psilocybin may instead have converged on a generic significance, such as awe, that any stimulus can take on. The interpretation most consistent with the network’s anatomy and item-level results is that modulation of experience by personal semantic and autobiographical history was weaker in participants reporting greater ego dissolution, so their representations in this network stayed closer to a response shared across people.

### Resting-state connectivity reconfiguration

Conversational partners, however, also converge on shared descriptions of ambiguous tangram figures (Clark and Wilkes-Gibbs, 1986) without losing their sense of self, so reduced idiosyncrasy alone is unlikely to be sufficient for ego dissolution. Because those partners still name what they see, a second condition may be that language-associated systems decouple from sensory input, so that perception loses the categories language ordinarily imposes on it. Connectivity scores for four other clades also correlated with EDI but, unlike c5’s (ρ=.455), did not survive correction (ρ=.313–.416).

The increase in c5’s thalamic coupling is consistent with two accounts. In Skipper’s (2022) predictive meta-analysis, thalamic activation overlapped speech-production maps but anticorrelated with the distributed language map and autobiographical-memory regions, which we interpreted as thalamic engagement in inner speech while its semantic periphery was inactive; the present thalamic increase and visual decrease have the same sign.

In the second, the cortico-striato-thalamo-cortical model, psychedelics reduce striatal inhibition of the thalamus and so increase thalamocortical transmission of less-filtered sensory and interoceptive information (Vollenweider and Geyer, 2001). Consistent with this model, ventral-striatal-to-thalamic effective connectivity decreases under LSD (Preller et al., 2019), thalamocortical coupling with transmodal targets increases (Avram et al., 2024) and, under LSD, dorsolateral prefrontal–thalamic connectivity is stronger in participants reporting greater ego dissolution (Coleman et al., 2025). Testing whether thalamic input drives the increase observed here requires directed measures such as effective connectivity.

Participants reporting greater ego dissolution simultaneously showed larger decreases in c5’s connectivity with posterior visual cortex (ρ=™.338, uncorrected p=.033; Figure 3A.iii), so this decrease may be an instance of the second condition. Alternatively, it may reflect direct drug effects on visual cortex: under LSD, participants whose primary visual cortex connected more widely reported stronger visual hallucinations (Carhart-Harris et al., 2016).

The connectivity score correlated most strongly with the four EDI items describing loss of ego and identity (ρ=.428–.512, Holm-adjusted p≤.034; Figure 3A.ii) and less with unity, reduced self-importance and reduced absorption in personal concerns (ρ=.219–.334). This division parallels recent scale development separating ego loss from unity (Sleight et al., 2023) and matches the hypothesis that language contributes more to reflective, narratively continuous identity than to oceanic unity or loss of self–world boundaries.

### Clinical analogues and testable predictions

Three of our observations parallel aphasia. First, psilocybin altered the connectivity of the left perisylvian network whose damage is strongly associated with aphasia, a disorder in which disconnection among surviving regions contributes to impairment (Catani and ffytche, 2005) and predicts it beyond lesion volume (Bonilha et al., 2014; Fridriksson et al., 2018). Second, the association was specific to loss of ego and identity, which aphasic stroke also disrupts (Shadden, 2005; Musser et al., 2015). Third, the more ego dissolution participants reported, the more c5 decoupled from posterior visual cortex, whose disconnection from language cortex causes pure alexia (Epelbaum et al., 2008) and, on one account, optic aphasia (Coslett and Saffran, 1989), and the more it coupled with the thalamus, whose damage is associated with thalamic aphasia (Crosson, 2013). We propose that psilocybin disrupts both pathways and that ego dissolution resembles a partial, reversible aphasia.

In pure alexia and optic aphasia, patients cannot read or name what they see, yet recognise it and can name it through other modalities (Coslett and Saffran, 1989; Epelbaum et al., 2008). Under LSD, picture naming stays fast but produces more semantically related substitutions (Family et al., 2016), so c5’s decoupling from visual cortex may be a partial counterpart. If so, psilocybin should impair naming from vision more than naming through other modalities, in proportion to ego dissolution.

Thalamic aphasia impairs lexical retrieval, spares repetition and co-occurs with disrupted prefrontal and frontotemporal networks (Crosson, 2013, 2021; Nishio et al., 2014; Rangus et al., 2021; Stockert et al., 2023), and on Crosson’s account the thalamus, under frontal control, selectively engages the cortex needed to retrieve a word. Psilocybin increased c5’s coupling with this thalamic and frontal circuit, and it broadens semantic activation more generally, as shown by increased indirect semantic priming (Spitzer et al., 1996). Looser lexical selection would produce this broader activation, and lexical selection is the function thalamic aphasia impairs, so drug and damage may disrupt the same process from opposite directions, one by increasing thalamic input and the other by removing it. Because naming deficits after thalamic damage in chronic left-hemisphere stroke were explained by accompanying damage elsewhere (Zhang et al., 2026), thalamic effects on naming may also be short-lived. If psilocybin loosens lexical selection, semantically related substitutions should increase with ego dissolution and with the change in c5’s thalamic coupling.

C5’s left lateralisation is also consistent with Levin’s (2025) proposal that psychedelics attenuate a normally dominant, language-mediated left-hemispheric mode, releasing a right-hemispheric mode characterised by broader contextual integration and weaker categorical parsing. Experiments involving unilateral intracarotid anaesthesia, which reversibly inactivates one hemisphere, provide evidence for this account: left-sided injection produces greater unawareness of comprehension deficits than right-sided injection, and loss of consciousness follows injection of a hemisphere in proportion to its language dominance (Serafetinides et al., 1965; Banks et al., 2010). In principle, intracarotid administration of a tryptamine such as DMT, whose effects begin and resolve within minutes, could permit an analogous test, despite considerable safety, pharmacokinetic and ethical barriers. We would predict greater ego dissolution after left-than right-hemispheric administration at matched doses, scaled by individual language lateralisation.

More practicable tests should target inner speech, because its loss would compromise the symbolic access through which experience is inspected, evaluated and assimilated within a temporally extended self-model (Skipper, 2022). In aphasia, inner speech can survive severe impairment of overt production, and people who retain it show better lexical retrieval and self-monitoring (Fama et al., 2019; Fama and Turkeltaub, 2020). Psychedelics preserve but reorganise language at the moderate doses used in priming and naming studies (Spitzer et al., 1996; Family et al., 2016), whereas higher doses can produce mutism and withdrawal from verbal exchange (Malitz et al., 1960). Because overt speech may be least available in the states of greatest interest, null effects on overt measures would be difficult to distinguish from a behavioural floor. Experience sampling across the acute window can already assess inner speech, and advances in non-invasive decoding of intended language make its continuous measurement increasingly conceivable (Lévy et al., 2026). Combined with individually localised language tasks under placebo-controlled psilocybin and narrative stimuli acoustically matched to unintelligible speech, these approaches would let researchers test whether disruption of inner speech tracks loss of ego and identity and whether the present association reflects language itself or another process with a similar anatomical distribution.

### Limitations

Inferring language involvement from spatial correspondence with meta-analytic maps is a reverse inference whose strength depends on how selectively the regions involved serve language (Poldrack, 2006). In defence of our approach, we nominated language in advance, so the inference was a falsifiable test: we predicted convergence centred on inferior-frontal and temporal cortex, and the clade that showed it, left-lateralised, transmodal and separate from the auditory and executive clades, matched the language map more strongly than any comparator clade. Direct measures of language and inner speech are needed to determine what the clade computed and the direction of any linguistic change.

Altered sensory processing may contribute: the psychedelic DOI impairs auditory deviance detection in mice, while psilocybin alters visually evoked responses in humans and effective connectivity within visual pathways (Kometer et al., 2013; Horrocks et al., 2025; Stoliker et al., 2025) and, in rats, lowers glucose metabolism in auditory and visual cortex one week after administration (Gudmundsen et al., 2026). Because c5 corresponded negligibly with the auditory and visual maps (r=.018 and r=™.059; Supplementary Data S2) and followed a different time course from the auditory (c4), visual (c2) and sensorimotor (c1) clades that also converged (Figure 2A.iii), a purely sensory account is unlikely to explain its convergence. Testing whether sensory changes disrupted linguistic organisation, or the reverse, requires manipulating one while measuring the other.

Because only the EDI items describing loss of ego and identity correlated with the connectivity score after correction, the claim is restricted to reflective and narrative selfhood (Millière, 2020). The EDI, completed once after scanning, measures a later reconstruction of the experience; that reconstruction is possible if minimal subjectivity and memory persist through compromised first-personhood (Metzinger, 2020; Fink, 2020). Elementary imagery was rated at the end of the day, after participants had returned to baseline, so it is a less immediate comparator than the EDI. The exploratory resting-state association survived correction across clades, but because the same participants also defined the target regions, it requires independent replication. Because applying a between-participant association to individuals assumes ergodicity, which human data often violate (Fisher et al., 2018), testing whether the association holds within individuals, and whether changes in language-associated systems drive, result from or accompany ego dissolution, requires ratings repeated during scanning or by experience sampling, ideally across doses, drugs and populations.

Only the directional joint-rank model was fitted, so the result is conditional on the hypothesis that high scorers converge while low scorers remain idiosyncratic. Component masks were selected for their strongest positive association without repeating threshold selection within each permutation, so component-level p values are conditional and mildly anti-conservative; because every analysis shares this bias, the analyses remain comparable. The fixed dendrogram cut prevented optimisation for the primary result, although language decoding varied with resolution. Because we hyperaligned each session separately, differences between sessions combine changes in representation with differences between the two alignment spaces. Sample sizes were modest for a multistage analysis.

## Conclusions

Under psilocybin, participants reporting stronger disruption of self-experience had more similar brain activity to one another in a distributed, left-lateralised network resembling the brain’s language system. At rest, this network became less connected with visual cortex and more connected with thalamic and frontal regions, especially in participants reporting greater loss of ego and identity. These findings are consistent with a transient disruption of language-mediated self-organisation as one contributor to ego dissolution. On this account, psilocybin reduces the influence of each person’s semantic and autobiographical context on present experience by altering language-associated systems and their connectivity with sensory regions. This account motivates experiments that manipulate and measure language, including inner speech, under psychedelics to establish its mechanistic role in maintaining a sense of self.

## Data and code availability

The source PsiConnect dataset is described by Novelli et al. (2026) and is subject to the access and participant-data-sharing terms specified by the source investigators. Code for the present analysis will be made available upon request.

## Author contributions

Author contributions should be finalised using the CRediT taxonomy. Provisional statement: Greg Cooper: Conceptualisation, Methodology, Software, Formal analysis, Writing - original draft, Visualisation. Vincent Sebastian Astl Delgado: Conceptualisation, Formal analysis of an earlier analysis version. George Blackburne: Conceptualisation. Devon Stoliker, Leonardo Novelli and Adeel Razi: Resources, Investigation and source-study data collection or preprocessing, Writing - review and editing. Jeremy Skipper: Conceptualisation, Supervision, Writing - review and editing. David Green: Conceptualisation, Writing - review and editing. Ravi Das: Supervision, Writing - review and editing.

## Funding

Greg Cooper, Ravi Das and Jeremy Skipper are supported by Wellcome Leap as part of the Untangling Addiction Program.

## Competing interests

GC – no competing interests

## Acknowledgements

We thank the PsiConnect participants and the wider source-study team. We acknowledge the facilities and scientific and technical assistance of the National Imaging Facility, a National Collaborative Research Infrastructure Strategy capability at Monash Biomedical Imaging, a Technology Research Platform at Monash University. We acknowledge and thank Leor Roseman for access to, and discussion of, the earlier LSD dataset that informed pilot analyses preceding the present psilocybin study.

## Supplementary Information

### Supplementary data tables

**Supplementary Table S1.** Dominant anatomical distribution and exploratory complete-vocabulary decoding of each clade. Anatomical descriptions use the complete positive-GMM maps. Terms are the highest-ranked cognitive/function terms surviving within-clade correction at q<.05; “None” indicates that no term survived. Full anatomical overlaps and term-level statistics are provided in Supplementary Data S1 and S3.

| Analysis | Clade | Dominant anatomy | Highest-ranked complete-vocabulary terms surviving $q < .05$ |
| --- | --- | --- | --- |
| **Psilocybin–EDI** | c1 | Right-dominant sensorimotor and dorsal-attention cortex, with default, salience/ventral-attention and control involvement | None |
|  | c2 | Bilateral visual cortex with dorsal-attention and default involvement, together with thalamic and cerebellar support | Visual; vision; motion; spatial; perceptual |
|  | c3 | Right-dominant lateral prefrontal control cortex, with default and salience/ventral-attention involvement | Executive; cognitive; cognitive control; executive function |
|  | c4 | Bilateral temporal–opercular and somatomotor cortex, with salience/ventral-attention and dorsal-attention involvement | Sound; auditory; tactile; pitch; touch |
|  | c5 | Left-lateralised distributed frontotemporal map spanning default, control, salience/ventral-attention and dorsal-attention territories | Syntactic; verb; language; sentences; semantic |
|  | c6 | Medial-temporal, limbic and default territories, with bilateral hippocampal, caudate, thalamic and cerebellar involvement | Stress; anxiety; reactivity; memories; emotional |
| **Psilocybin–SI** | c1 | Bilateral prefrontal default and control cortex | Prospective; choice |
|  | c2 | Compact, strongly right-lateralised amygdala–hippocampal, orbitotemporal and striatal map | Fearful; fearful faces; neutral faces; fear; neutral |
|  | c3 | Strongly left-lateralised premotor, dorsal-attention and frontoparietal-control cortex | Tasks; planning; motor imagery; perform; complexity |
|  | c4 | Bilateral temporal–opercular, somatomotor and salience/ventral-attention cortex | Pitch; auditory; sounds; sound; acoustic |
|  | c5 | Bilateral occipital visual and dorsal-attention cortex | Motion; visual; visual motion; vision; visual stimuli |
|  | c6 | Right-lateralised frontoparietal control and dorsal-attention cortex | Task difficulty |
| **Baseline–EDI** | c1 | Bilateral frontal default and control cortex | None |
|  | c2 | Compact occipital visual map | None |
|  | c3 | Sensorimotor cortex with dorsal-attention and salience/ventral-attention involvement | Sensorimotor; motor task; motor; movement; motor imagery |
|  | c4 | Posterior-medial default, visual and dorsal-attention cortex, including precuneus and posterior cingulate territories | None |
|  | c5 | Strongly left-lateralised temporal–opercular, somatomotor and salience/ventral-attention cortex | Sound; auditory; sounds; tones; pitch |
|  | c6 | Large distributed map spanning default, visual, salience/ventral-attention, control and limbic territories, with hippocampal, thalamic, striatal and cerebellar involvement | None |
| **Baseline–SI** | c1 | Bilateral frontal default and control cortex | Executive; rules |
|  | c2 | Bilateral occipital visual cortex with substantial cerebellar support | Visual; object recognition; motion; visual motion; visual perception |
|  | c3 | Large distributed map spanning default, visual, salience/ventral-attention, somatomotor and control territories, with hippocampal, striatal and cerebellar involvement | None |
|  | c4 | Strongly left-lateralised temporal–opercular and somatomotor cortex | Auditory; sound; sounds; tones; pitch |
|  | c5 | Posterior parietal map spanning dorsal-attention, default and somatomotor territories | None |
|  | c6 | Right-dominant premotor, control, dorsal-attention and salience/ventral-attention cortex | None |

**Supplementary Data S1**. Anatomical inventory of all 24 analysis-by-clade GMM-core maps, including voxel counts, Schaefer-1000 cortical parcel overlaps, subcortical overlaps and Figure 2 display ranges (Supplementar yData_S1_clade_anatomy_summary.csv; SupplementaryData_S1_clade_anatomy_overlaps_long.csv).

The complete positive-GMM clade maps varied substantially in size and anatomical distribution (489–13,364 voxels; Supplementary Data S1). Within psilocybin–EDI, c1 was weighted toward somatomotor and dorsal-attention territories, c2 toward visual cortex, c3 toward frontoparietal control and default-network cortex, c4 toward somatomotor, opercular and superior-temporal territories, and c6 included hippocampal, limbic and medial-posterior regions. The language-associated c5 map comprised five significant ICs and 5,701 voxels. It was distributed but left-lateralised, peaking in dorsal pars opercularis of the left inferior frontal gyrus (MNI [™45, 16, 30]) and extending across temporal, lateral-prefrontal and frontal-opercular/insular territories in default, control and salience/ventral-attention networks. Maps in the other three analyses showed similarly distributed but distinct anatomical configurations. These summaries use the complete GMM-core maps rather than the 75th-percentile masks used for display.

**Supplementary Data S2**. Complete a priori Neurosynth results for all four analyses, six clades and 14 terms (neurosynth_apriori_complete_4cells_6clades_14terms.csv). Twelve of the 336 a priori clade-by-term tests survived correction at q<.01 (Supplementary Data S2). Auditory correspondence was present in every analysis, although the winning clade varied: psilocybin–EDI c4 (r=.381, q=.0028), psilocybin–SI c4 (r=.458, q=.0017), baseline–EDI c5 (r=.415, q=.0084) and baseline–SI c4 (r=.456, q=.0042). Visual correspondence was significant for psilocybin–EDI c2 (r=.353, q=.0028), psilocybin–SI c5 (r=.540, q=.0017) and baseline–SI c2 (r=.322, q=.0042), but not baseline–EDI. Language-map correspondence met q<.01 only for psilocybin–EDI c5 (r=.382, q=.0028). Speech-perception correspondence was also observed for psilocybin–EDI c4 and psilocybin–SI c4, while speech production and interoception were significant only for psilocybin–SI c4.

### Exploratory complete-vocabulary Neurosynth decoding

*Supplementary Data S3. Five highest-ranked complete-vocabulary cognitive/function terms for every analysis and clade after documented anatomical, methodological, population and clinical/index exclusions (neurosynth_complete_vocabulary_top5_cognitive_by_cell_clade*.*csv)*.

Exploratory complete-vocabulary decoding further differentiated the psilocybin–EDI clades (Supplementary Data S3). C2 was characterised by visual, motion and perceptual terms; c3 by executive and cognitive-control terms; c4 by sound, auditory, pitch and tactile terms; c5 by syntactic, verb, language, sentences and semantic terms; and c6 by stress, anxiety, reactivity, memory and emotion terms. C1 had no term surviving complete-vocabulary correction. Psilocybin–SI instead separated fear- and face-related c2, planning and motor-imagery c3, auditory c4 and visual-motion c5. At baseline, corrected terms were more restricted: baseline–EDI showed sensorimotor decoding in c3 and auditory decoding in c5, whereas baseline–SI showed executive/rule-related c1, visual c2 and auditory c4.

### Supplementary figures

The direction of psilocybin’s effect on stimulus-locked synchrony depended on the task (Supplementary Figure 1A–B). Mean whole-brain ISC decreased during music, from r=.1007 at baseline to r=.0890 under psilocybin, but increased during cloud viewing, from r=.0860 to r=.0947. Although the autocorrelation of the spatially averaged ISC trajectory crossed zero later under psilocybin in both tasks, this did not generalise to local voxelwise trajectories: the median local crossing was slightly earlier during music and unchanged during clouds. In participant-level leave-one-out analyses, local synchrony persisted for less time under psilocybin during music (Supplementary Figure 1C). Mean first-zero crossing decreased from 7.14 to 5.56 s, while positive autocorrelation-function (ACF) area over 0–30 s decreased from 1.39 to 1.19 correlation·s. After two-sided voxelwise FWER correction and the >20-voxel descriptive reporting filter, the effects comprised 175 and 1,888 voxels, respectively, and were uniformly negative. The largest ACF-area reductions involved bilateral auditory/perisylvian and temporoparietal cortex, medial occipital and precuneus cortex, with smaller frontal, paracingulate and lateral-occipital components. No cloud-viewing component exceeded 20 voxels after voxelwise correction.

**Supplementary Figure 1.**
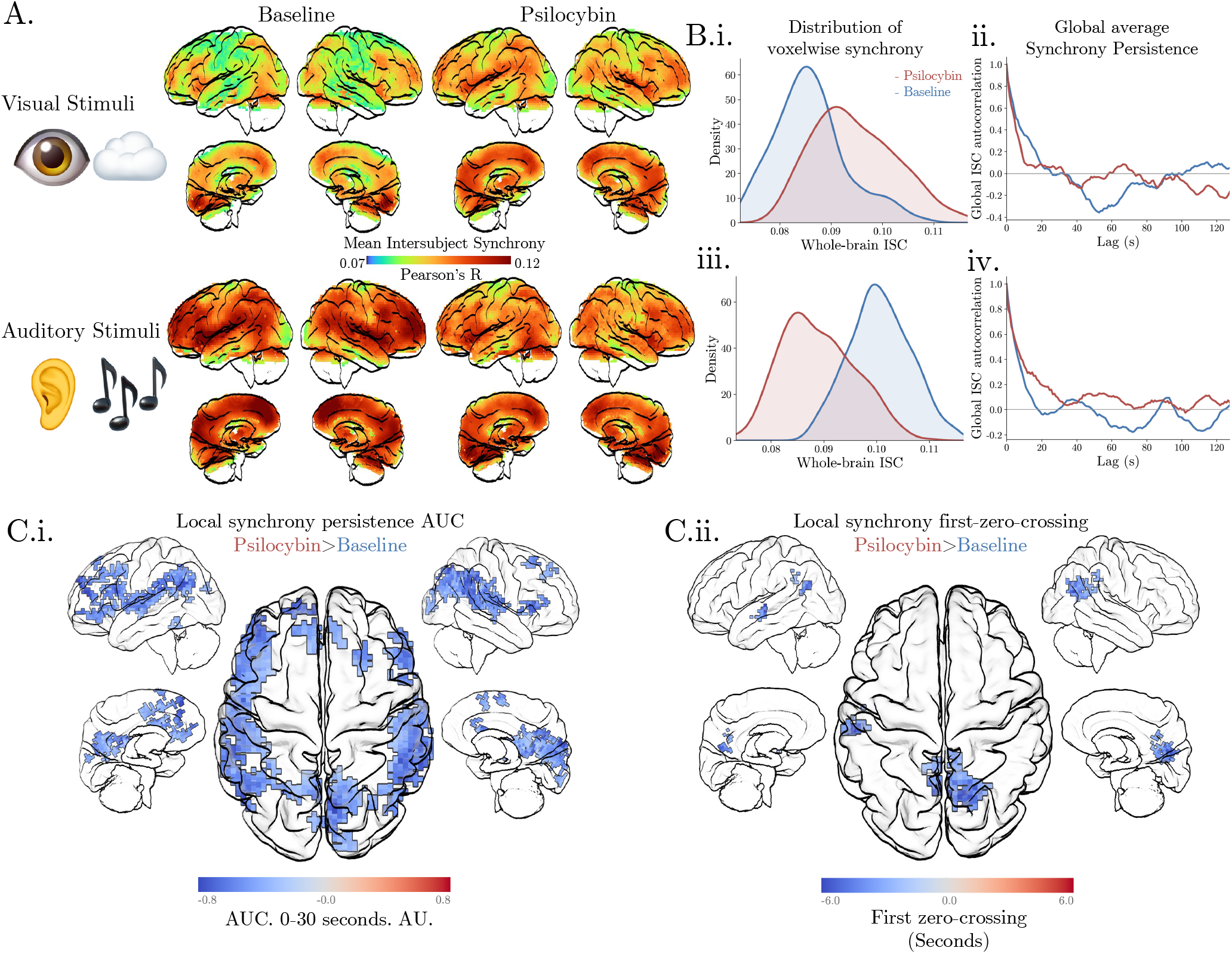
Descriptive inter-participant synchrony. (A) Mean ISC maps at baseline and under psilocybin during cloud viewing (top) and music (bottom). (B) Distributions of voxelwise ISC (i, iii) and autocorrelation of the spatially averaged ISC trajectory (ii, iv) during cloud viewing (i–ii) and music (iii–iv). (C) Voxelwise paired psilocybin-minus-baseline contrasts of how long local synchrony persisted during music: (i) positive autocorrelation area over 0–30 s and (ii) first zero crossing (two-sided voxelwise FWER correction; clusters >20 voxels); no cloud-viewing component survived.

## Notes

### Competing Interest Statement

The authors have declared no competing interest.

